# Biocultural vulnerability of traditional crops in the Indian Trans Himalaya

**DOI:** 10.1101/2024.10.22.619747

**Authors:** Harman Jaggi, Akshata Anand, Katherine A. Solari, Alejandra Echeverri, Rinchen Tobge, Tanzin Tsewang, Kulbhushansingh Suryawanshi, Shripad Tuljapurkar

## Abstract

Traditional agricultural landscapes are critical for conserving biocultural and ecological diversity. Despite their significance, traditional systems have often been overlooked, leading to genetic erosion of crop landraces. Using the fragile ecosystems of northwest Himalaya as case study, we examine the ecological and genetic resilience of an understudied and lesser-known traditional crop, black pea, and barley (*Hordeum vulgare*) and compare them to the introduced cash crop of green pea (*Pisum sativum L*.). We co-designed field experiments with local farmers to assess survival and reproductive traits for the crops. We performed whole-genome sequencing to investigate the genetic diversity of black pea and describe their nutritional profile. Our findings indicate that traditional crops are better adapted to local climatic conditions and hold considerable genetic diversity and nutritional potential. We emphasize the importance of integrating traditional knowledge with scientific research to promote sustainable food systems and socio-ecological stability in vulnerable mountain regions.

## 1 Introduction

Traditional farming landscapes and mountain communities are important reservoirs for conservation of biodiversity, biocultural diversity and agro-ecological resources [15, 49, 16, 45, 19]. Traditional farming knowledge (in the form of, e.g., local crop landraces and seedbanks) results from complex cultural and environmental selection [2] and supports diverse ecosystem services [22, 9]. However, traditional practices have often been overlooked in conservation and modern agronomic research [3] even as researchers have noted declines in the diversity of landraces and crop genetic erosion [26]. This is concerning because traditonal production systems (*e*.*g* locally adapted crop varieties) may inform nutritional, cultural and ecological stability, especially in adverse climates [51, 7]. Therefore, research on local production systems is likely important for supporting regional livelihoods, sustaining socio-ecological stability and preserving cultural knowledge [58, 3].

The stability and productivity of food systems is increasingly vulnerable to increased variation in weather patterns, and in the frequency of extreme events (such as episodes of drought) often due to climate change [5, 32]. The high-altitude Trans-Himalayan landscape presents excellent case studies for examining traditional food systems and the impacts of climate change on agricultural production in mountainous communities. In that region, and more broadly over Asian montane rangelands, climate is changing even faster than the global average [10, 27], threatening both the agroecological landscape and local economy. These rangelands provide exceptional biodiversity and cultural richness [37, 54, 41], and are also known as the Third Pole, because they hold the next largest reserves of freshwater and glacial ice after the polar regions. The human communities in these cold desert ecosystems share habitat with diverse wildlife, including the iconic but vulnerable snow leopard *Panthera uncia*, and depend closely on ecosystem services.

Our research examines traditional agricultural systems and their vulnerability in the fragile ecosystems of northwest Himalaya, in the state of Himachal Pradesh, India in Spiti district (map 1). Agricultural production in Spiti was traditionally subsistence-based [33]. The main crops grown were barley *Horedum vulgare* (locally *neh* or (*jau*)), and a local variety of pea called black pea (*sanmoh nako* or *dhoopchum* in Tibetan). The scientific name for black pea remains unclear (as we later explain) and until recently [50, 6] there were no scientific research articles on black peas. In 1983, an agro-economic revolution in Spiti was spearheaded by a family that experimentally planted green pea or garden peas *Pisum sativum L*. [33]. In the subsequent two decades, the predominantly subsistence-based production of the village communities has been increasingly replaced by the market-oriented production of cash crops (mainly green peas, apples), destabilizing the sustainability of the cropping system [44, 35]. During our socio-cultural work (*manuscript in prep*), we learnt that black pea and barley are an integral part of ethnodietary recipes and of a climate-resilient farming system, but their cultivation has drastically declined due to market forces and lack of information. The practice of polyvarietal cultivation of barley (*kneu, soha, nenak, and eumo*) is nearly extinct [33]. *kneu* is the only variety grown currently with small quantities of *soha* for religious ceremonies. Without adequate research, the understudied black peas are close to facing cultural extinction.

Grown for many centuries, barley and black peas are likely well adapted to the region’s geoclimatic conditions and irrigation regimes. Here, we provide an ecological, genetic and nutritional characterization of black pea, which to our knowledge has not been done before. We compared several aspects of traditional crops (we include black peas and barley) to the cash crops of green peas. The key research questions are as follows: firstly, we examined whether traditional crops are better adapted to changing climatic conditions in terms of their survival and reproductive traits. Secondly, we examined whether black pea differs in genetic diversity from other peas in the *Pisus* genus. Thirdly, we examined the nutritional profile of back pea.

To examine our first question, we performed field experiments (March-September 2023) to compare the three different crops (black pea, barley and green pea) for reproductive and photosynthetic traits under variable water treatments across three elevation gradients (see Materials and Methods). We co-designed the experiments with local farmers who advised on sowing depth and bed preparation to ensure optimal growth conditions. We hypothesized that traditional crops are more resilient than cash crops as measured by survival and reproductive traits.

To examine the second question, we performed whole-genome sequencing (shotgun sequencing using the Illumina platform) for a black pea sample (see Materials and Methods). We then used a chromosome-wide Principal Component Analysis (PCA) to compare our black pea sample with pea accessions published in [59]. We expected genetic differences between black peas and green peas, but did not have a prior on the magnitude and nature of those differences. For the third question, we collaborated with the Central Food Technological Research Institute (CFTRI) in India to provide a standardized nutritional profile for black peas. Nutritionally, we expected that black peas have a distinct and high nutritional profile in comparison to green pea because cash crops are consistently selected for particular nutritional values and sold in markets for that trait [47].

Our findings suggest that black peas and barley are not only better adapted to local conditions (with higher survival, flowering and pod production) but that black pea also holds significant untapped genetic and nutritional potential. Our research aligns with a growing body of evidence that advocates for a reevaluation of traditional and Indigenous agricultural practices [3]. In the next sections, we describe the study system, research methods, results and recommendations. We close with a discussion on integrating local knowledge and modern science to examine lesser-known and under-utilized crops to foster food systems that are environmentally sustainable and adaptable to changing climatic conditions.

## 2 Results: Traditional production systems

### 2.1 Results from field experiments

We first want to note that the results are averaged across water treatments because of high rainfall in the summer of 2023. The drought treatments received rainfall and we therefore treated different treatments as replicates. Our first result from the field experiment shows both black peas and barley have higher survival probability (or establishment rate since few seeds are eaten by birds, livestock and ungulates) in comparison with green peas across all sites. The results are consistent across three different ecological settings as shown in Figure 2. The site Kiamo (at 4000m) has highest average establishment rate for black peas with 0.51 overall. The high establishment rates for black peas (evidenced by the higher mean value) underscore a possible genetic or phenotypic advantage over green peas. These findings suggest that black peas may possess attributes that confer greater survival advantages under the diverse conditions represented by the study sites.

**Figure 1.**
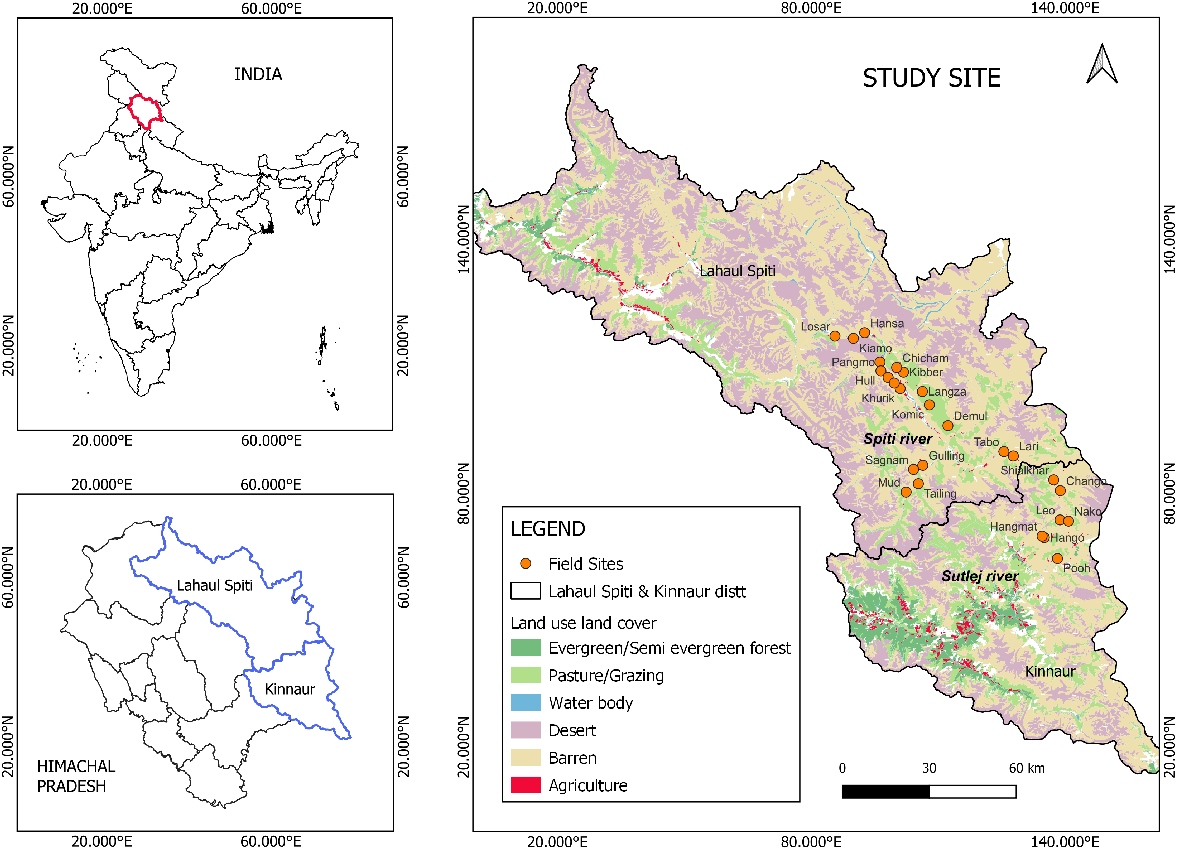
The right panel shows location of the sites within Spiti Valley and Kinnaur district where we interviewed local community members across 26 villages and helped co-design experimental plots at the three sites discussed in the Methods section. Different colors correspond to land use characterizations such as agriculture, grazing pastures, and barren among others. The orange solid circles correspond to villages sampled. The bottom-left panel shows the extent of Spiti and Kinnaur district within the state of Himachal Pradesh, India.

**Figure 2.**
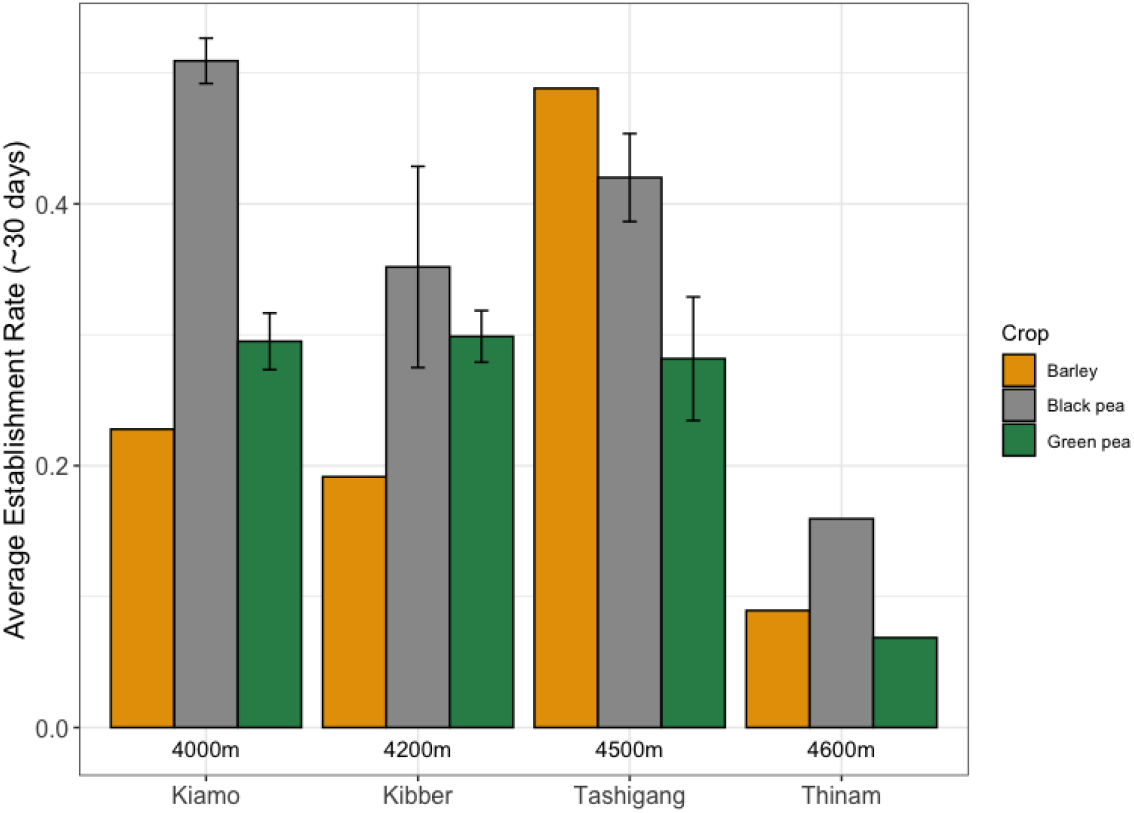
Establishment rate (survival probability) of black peas, green peas, and barley after 30 days. The x-axis corresponds to four sites at different elevations: Kiamo (4000m), Kibber (4200m), Tashigang (4500m), and Thinam (4600m). The y-axis corresponds to average establishment rates for the crops. Each color corresponds to different crops: barley, black pea and green pea. The lack of error bars indicate there was one replicate.

**Figure 3.**
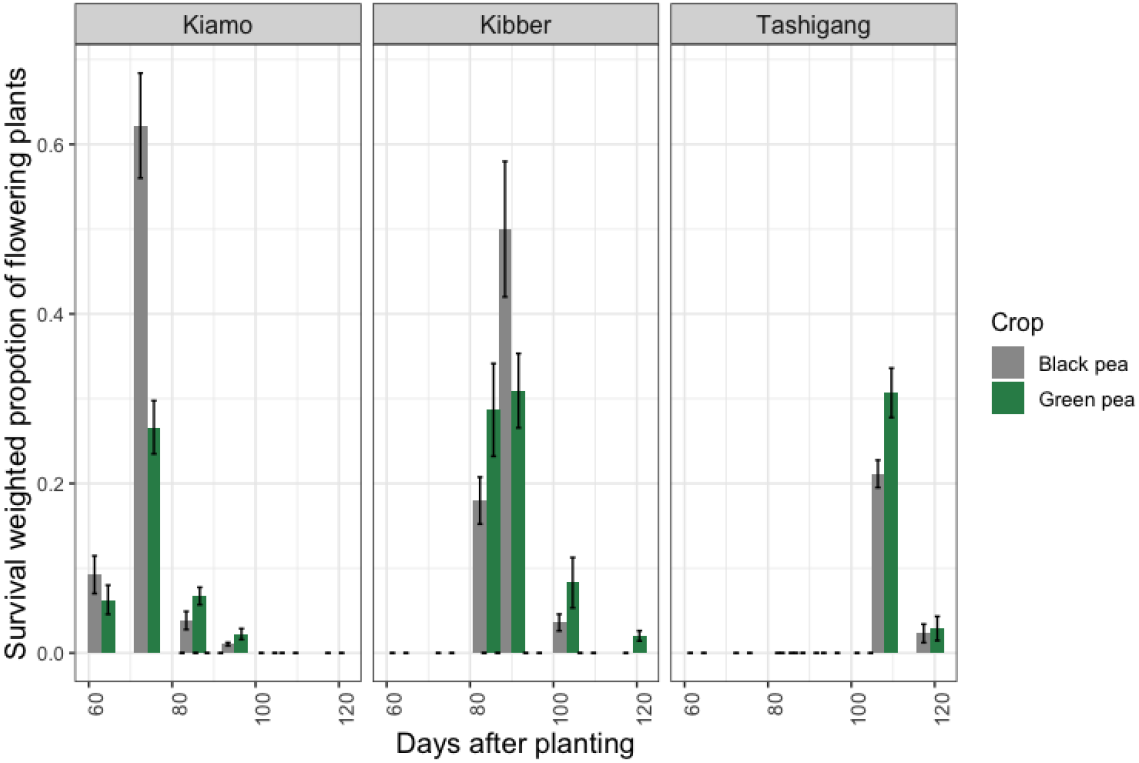
Survival-weighted proportion of flowering plants. Each panel corresponds to three sites at different elevations: Kiamo (4000m), Kibber (4200m), Tashigang (4500m). The x-axis corresponds to days after planting the seeds and the y-axis corresponds to survival-weighted proportion of flowering plants. Each panel corresponds to three sites at different elevations: Kiamo (4000m), Kibber (4200m), Tashigang (4500m). The x-axis corresponds to days after planting the seeds and the y-axis corresponds to survival-weighted proportion of flowering plants. The color corresponds to black peas and green peas.

Except Tashigang, barley establishment rates are lower than green peas across all sites. Please note there are no error bars for barley because some of our barley plots were destroyed and we were only able to preserve one plot per site. We also added another experimental site Thinam at 4600m without replicates to note seed survival in high elevation. Farmers noted they had been experimenting growing crops at even higher elevation (4600m) due to better availability of snow melt and access to water. We find low success at this elevation across all crops at Thinam as shown in last panel of Figure 2.

Next, we evaluate non-reproductive traits such as average stem height and average number of leaves (correspond to higher photosynthetic activity). As shown in the Supplementary figures, black peas outperform green peas consistently across all sites. The average stem height for black peas is 38.12cm, 24.3cm, 25cm compared to 18.2cm, 12.48cm, 10.7cm, across Kiamo, Kibber, and Tashigang, respectively. We do not consider barley in these plots because barley is a grass species from *Poaceae* family.

Important covariates for farmers include the reproductive traits such as proportion of flowering plants and pea pods. Figure 3 describes the distribution of survival-weighted proportion of flowering plants. Through the cropping season, we noted the proportion of flowering plants and weighted it by survival. We find black peas have higher proportion of flowering plants for black peas than green peas. Similarly, the average number of pods per individual plant are also higher for black peas than green peas as shown in Figure 4. Finally, we evaluate the growth rate for all crops by taking stem stem height as a predictor and find black peas, barley and green peas have growth rates 0.83, 0.6, and 0.38 as shown in Figure 5.

**Figure 4.**
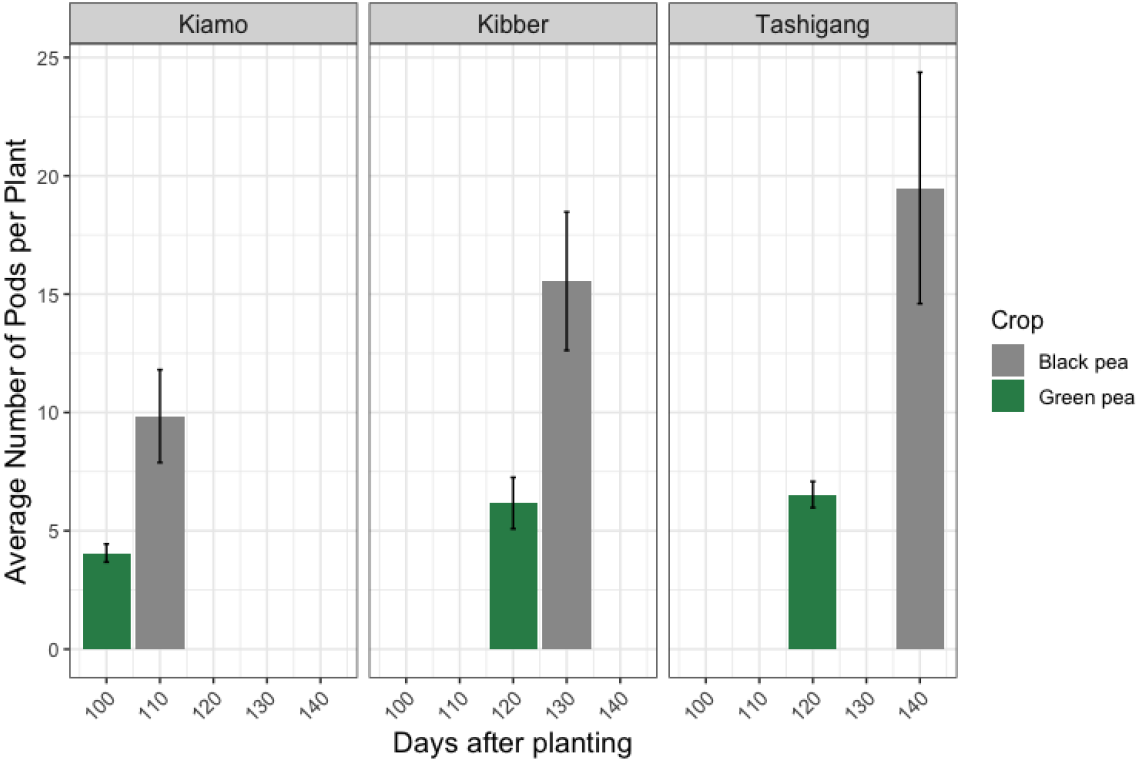
Number of pods per individual. Each panel corresponds to three sites at different elevations: Kiamo (4000m), Kibber (4200m), Tashigang (4500m). The x-axis corresponds to days after planting the seeds and the y-axis corresponds to number of pods per individual plant. The color corresponds to black peas and green peas.

**Figure 5.**
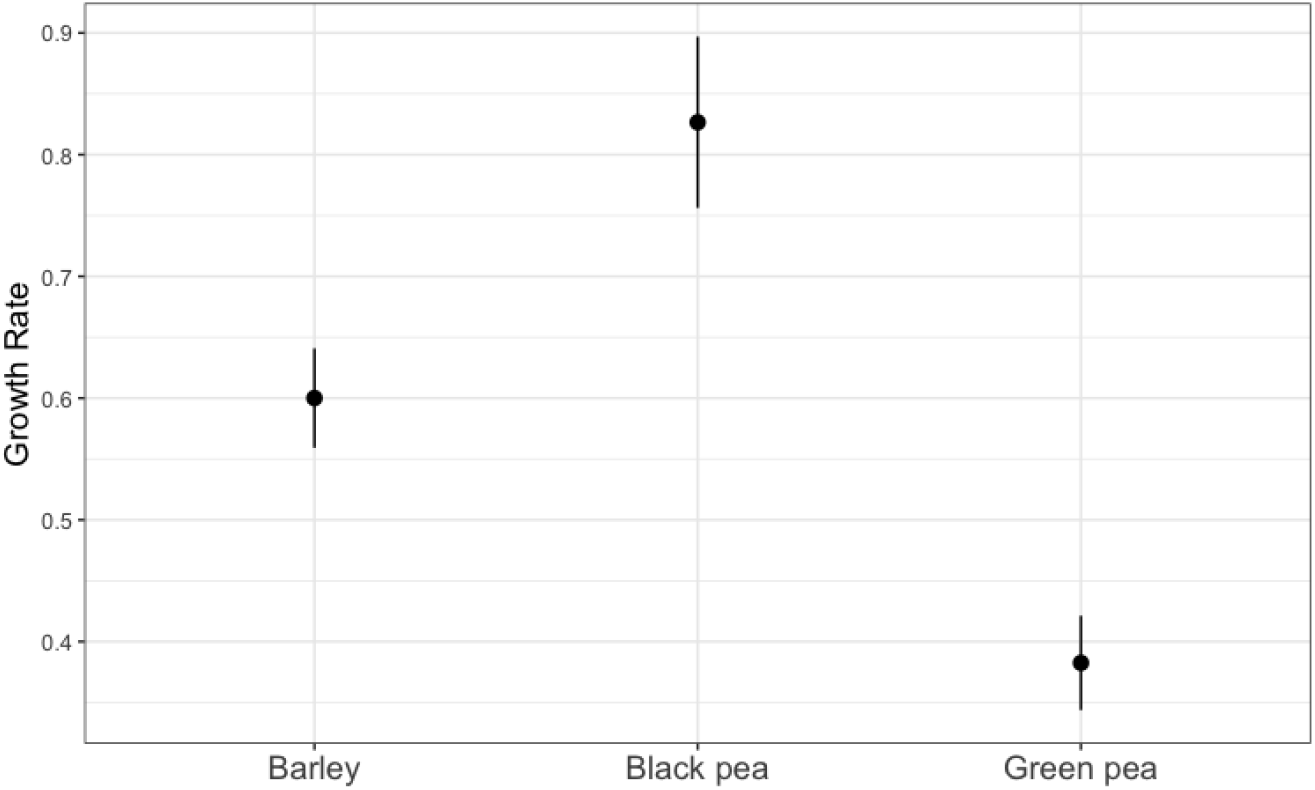
Growth rate of each crop: black peas, green peas, and barley for all sites. The y-axis corresponds to the log growth rate and x-axis corresponds to the three crops.

### 2.2 Genetic diversity of black peas

Pea belongs to the most ancient set of cultivated plants and is known to be domesticated about 10,000 years ago in the Middle East and Mediterranean region (also known as Fertile Crescent) [1, 52]. Although the taxonomy of *Pisum* genus is complex and still under debate [59, 46], there is consensus that the genus consists of two species-wild *P. fulvum* and *P. sativum* [39]. *P. sativum* includes wild (subsp. *elatius*) and domesticated taxa (subsp. *sativum*).

We have generated the first whole genome sequencing data for the black pea and have mapped it to the P. sativum reference genome. The average depth of coverage of our black pea data was 15X with a breadth of coverage across the reference genome of 86.6% . We combined this data with data for pea accessions in [59], and performed a chromosome wide-PCA analysis to infer the structure of Pisum collection. Since we were interested in our black pea subpopulation, we used samples from *P. sativum* and admixed accessions. In addition, we included *Pisum sativum*

*var. arvense* (Accession SRR19543725) and performed a PCA for each chromosome to compare the samples. PCA results reveal the three principal components (PCs) explain almost 33.3% of the observed genetic variation in the collection.

We find pea accessions were separated based on the *P. sativum* subspecies consistent with previous studies [23, 59]. However, our black pea sample clustered distinctly away from *P. sativum subsp. sativum var. arvense* sample. In addition for chromosome NC_066581 and NC_066585 (as shown in Figure 6) our sample clusters neither with domesticated varieties nor the wild germplasms. Although our morphological study of black pea plant supports two recent studies [6, 50] from Himalayan region that identify the local black pea to be *Pisum sativum var. arvense*, a winter legume belonging to the Fabaceae (Leguminosae) family, our genetic results question the claim. Our research emphasizes the high-altitude Himalayan mountains to be a relevant secondary centre of pea diversity, rich in primitive cultivated forms of field peas [23].

**Figure 6.**
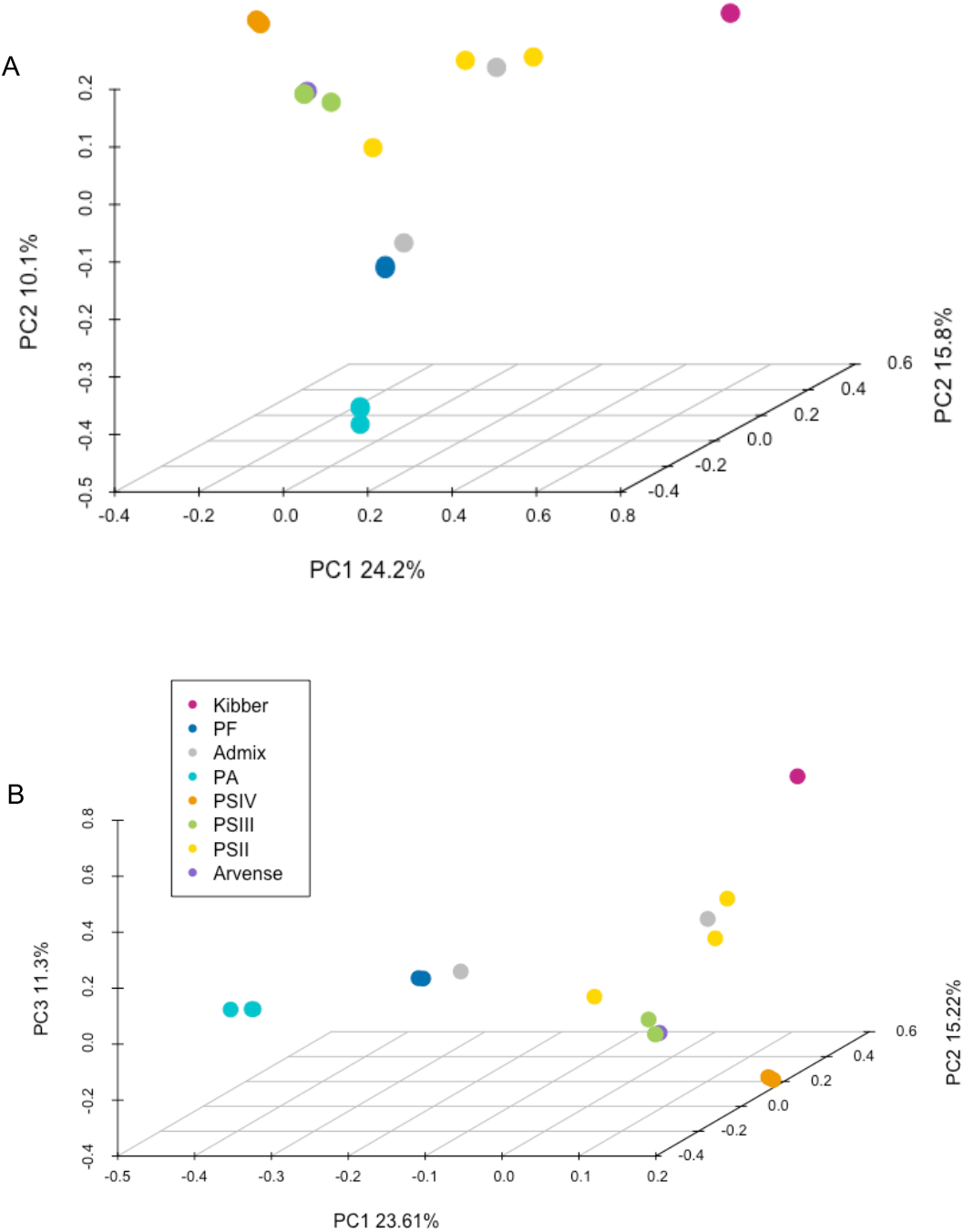
The figure shows PCA results for chromosome NC_066581 in panel A and NC_066585 in panel B. Kibber (in pink color) represents our sample and is distinct from *arvense* (in purple color) and other domesticated samples such as Pisum II and Pisum III (in green and yellow).

#### 2.2.1 Nutritional composition of black peas

Anecdotally, black peas are known for their diverse nutritional profile but the nutrient content for black peas from the Trans-Himalayas has been unclear. We collaborated with CFTRI based in Mysuru, Karnataka, India to create a nutritional profile for black peas. We find these legumes are rich in a variety of essential nutrients and offer a diverse combination of protein, fiber, vitamins, and minerals, making them not only culturally and ecologically important but also a valuable addition to a balanced diet. Table 1 provides a measure of nutrients and daily recommendation averages taken from NIH for a adult of body weight 70kg. Firstly, black peas are notably high in protein, with approximately 21.20% protein content per 100 grams. This makes them an excellent plant-based protein source, especially beneficial for individuals following vegetarian or vegan diets. Furthermore, black peas are a good source of dietary fiber (15.9% per 100 grams) in addition to crude fiber at 6.23% per 100 grams. Black peas also provide a range of essential vitamins such as Vitamin C (13.1 mg/100g), Vitamin B1 (0.61 mg/100g), Vitamin B3 (1.62 mg/100g). They are a good source of minerals such as iron (9.11 mg/100g), zinc (2.48 mg/100g), calcium (108.09 mg/100g), and magnesium (109.28 mg/100g).

**Table 1.**
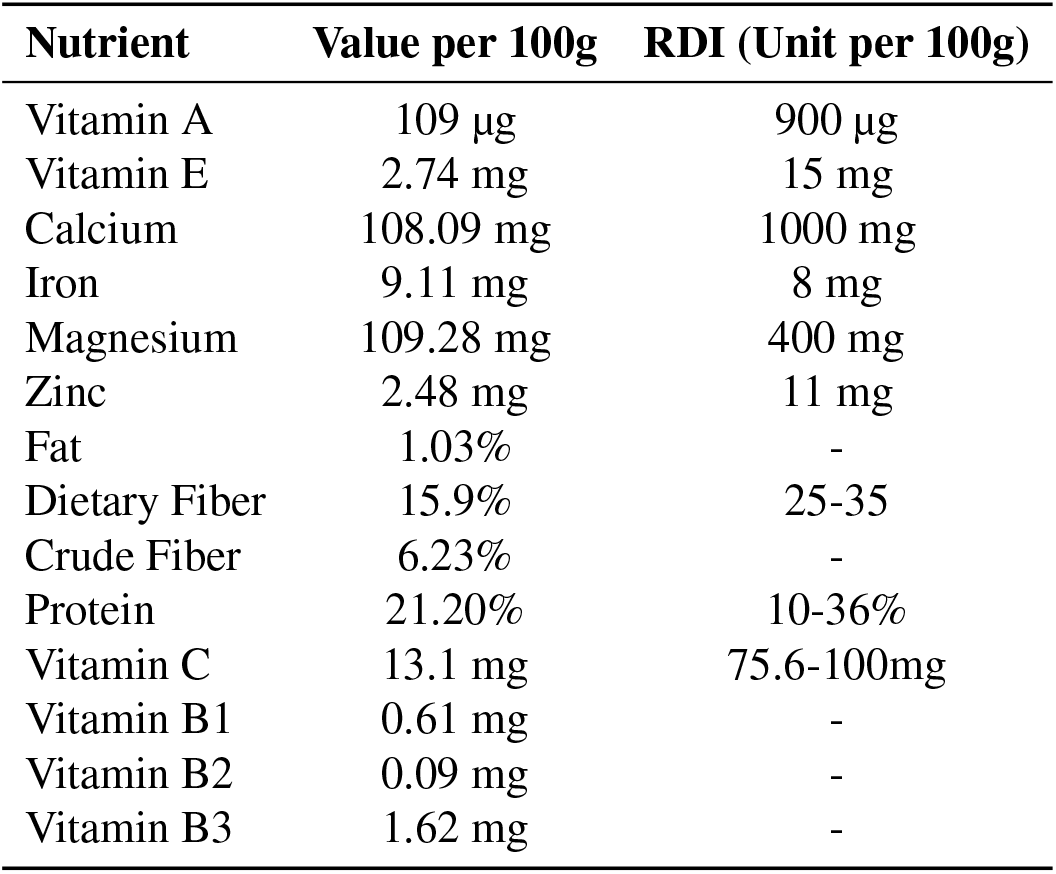
Nutritional Profile for black peas and Recommended Daily Intake (RDI) taken from National Institutes for Science (NIH) for an average adult of body weight 70 kg. The tests were performed at Central Food and Technological Research Institute, Mysuru, India. The % by wt corresponds to percentage of nutrient per 100 g.

## 3 Discussion

We find great potential for lesser-known and underutilized traditional crops such as black peas in terms of their survival and reproductive traits. We find black peas outperform green peas (cash crop) in terms of their establishment rates, average stem height and survival-weighted proportion of flowering plants (Figures 2, **??**, 3). The findings align with our socio-cultural research during which local farmers underscored black peas are better adapted and more resilient (in terms of ease of growing, and drought resistance) than green peas. Our multi-pronged approach provides a comprehensive baseline for traditional crops in high-altitude Trans-Himalaya as these food systems are crucial to the stability of humans and their ecosystems [12, 14, 42].

Further, we generated the first whole genome sequencing data for black peas from the Trans-Himalayan region to assess how genetically similar they are to other pea species. As shown in Figure 6 our sample clusters neither with domesticated varieties nor the wild germplasms. We expected genetic differences between black peas and green peas, but did not have a prior on the magnitude and nature of those differences. Although researchers identify black pea to be *P. sativum subsp. sativum var. arvense*, our findings show black peas to be distinct from *arvense*. Our research aligns with previous research [23] that emphasizes the high-altitude Himalayan mountains to be a relevant secondary centre of pea diversity, rich in primitive cultivated forms of field peas [23]. We also find black peas have high nutritional profile, offering high percentage of protein, dietary fiber, zinc, magnesium, among others. This makes them an excellent plant-based protein source, especially beneficial for individuals following vegetarian/vegan diets.

Locally inclusive development may enhance the sustainability of a traditional agricultural system, addressing both its conservation and its development. Our research emphasizes greater recognition of and research on lesser-known traditional crops in collaboration with communities who have long been using them, and who can be the main beneficiaries. Many studies have noted [29, 18] wild pea diversity in peripheral regions is insufficiently represented in the existing germplasm collections. [29] found two small populations of the wild pea *Pisum sativum subsp. elatius* in the Zagros Mts, Iran. They note wild pea populations are rare, small in size and suffer from climatic change and land exploitation. Another example is *Pisum sativum subsp. abyssinicum*, locally known as Dekoko peas, a unique subspecies endemic to Amhara and highland areas of Tigray, Ethiopia and to South Yemen. Dekoko peas are an important crop, widely mixed with the main cereal crops such as, sorghum, barley and teff [17, 18]. The crop qualifies for species status on the basis of its phenotype (early flowering and strongly serrate leaflets) and biological isolation [25]. Although cultivation of Dekoko peas had drastically declined due to drought and political disruption, in the past 10 years, there has been a considerable renewal of interest, resulting in new collections of Dekoko germplasm, characterization studies aimed at crop improvement and extending the areas of cultivation [55].

Looking beyond the *Pisum* genus, we can find many other examples of species may also be useful in addressing different effects of climate change. Thus, *Coffea stenophylla* can tolerate higher temperatures than the commercial species *Coffea arabica*, under threat from climate warming.

Communities in Sierra Leone helped researchers at the Royal Botanic Gardens (Kew) to find coffee berries from *Coffea stenophylla* in the wild [11]. In southwestern Ethiopia, the banana-like Enset (*Ensete ventricosum*)has starch-filled stems that are a main source of calories and nutrients, while the leaves are used for basket making, to feed cattle, and to provide shade and build roofs. Additionally, this species is remarkably tolerant of drought and short-term temperature variations [3]. And as a final example, we note that the UN recognized 2023 as the International Year of Millets [57], calling attention to their many nutritional and health benefits, their suitability for cultivation under adverse and changing climatic conditions, and their potential to contribute to the UN 2030 Agenda for Sustainable Development.

Environmental and conservation outcomes could improve by recognizing and honoring the relationships between people and their landscapes, and proposing actions based on local priorities [13, 34, 13]. Traditional agricultural systems are known for providing the communities with self-sustenance and sufficiency, while conserving the agroecosystems and associated biodiversity.

Globally and/or Nationally Important Agricultural Heritage Systems (GIAHS/NIAHS), identified by the Food and Agriculture Organization of the UN, recognize and designate bioculturally important sites to safeguard agrobiodiversity, protect agricultural heritage and recognise the knowledge and practices of local communities [28, 48, 14, 3]. These declarations have triggered the development of different tourism activities related to biocultural inheritance. In India the Indigenous ‘Bari System of Farming’ practiced by the native people of North Eastern India (Assam) is recognized from the perspectives of consumption, conservation, and biodiversity management [8]. The campesino cooperatives in Mapuche-Pewenche territory (southern Andes) practice community-based tourism incorporating Mapuche rural culture and agricultural practices that support local food systems and economy with a strong identity component [24]. We recommend the Trans-Himalaya for recognition as a GIAHS/NIAHS given the fragile ecosystems, high cultural and ecological diversity.

There are important limitations to out work. Most notably, we present findings from only one year of data collection. A consistent study over multiple years is needed to identify effects of climate change on the agroecosystem. The long-term datasets may also shed light on other demographic patterns-which stage in life-cycle of crop is most vulnerable to water stress and plant pathogens? What makes traditional crops more resilient? A quantitative understanding of demographic tradeoffs as well as effects of stochasticity on dispersal patterns will yield important insights for farmers and management interventions [56, 20, 21]. Further research is needed to identify the level of genetic variation and what genes and loci may be associated with adaptive traits [40]. This includes identification of desirable genes and traits from unadapted materials, such as CWRs, and their transformation for creating resilient cultivars [43].

## 4 Materials and Methods

### 4.1 Study site

In the state of Himachal Pradesh (HP), about 26,000 km^2^ area is potential snow leopard habitat [54]. Within this habitat, we focused on Lahaul-Spiti district (Figure 1). The area lies in the rain-shadow of the main Greater Himalayan range, within the Trans-Himalaya region, bordering Tibetan plateau in the east. The landscape is typically above the treeline, between 3000-6000 m elevation range. Temperatures range from -30^*°*^C in peak winter to 30^*°*^C in peak summer. The landscape is mountainous cold desert and rocky, with steep slopes largely dominated by grasses and shrubs. Precipitation is received primarily in the form of snow in between November and March, and starts to melt by April.

The major river system is the Spiti river that originates from Kunzum range and travels 150 km in Kinnaur district before its confluence at Khap with Satluj river (from Tibet). Spiti river flows in deep chasms and gorges for most of its length with a catchments area is about ∼ 12000km^2^ [53], most part being barren rocky land covered with thick moraines. The heavy snowfall in the winter months contributes to the Satluj’s flow in spring. The Pin is the largest tributary of the Spiti River and Lingti, Parechu are other important tributaries. The townships and settlements are clustered close to the river systems or near a glacier from which they harvest water.

Spiti valley has a low human population density (c. 1 person/km^2^). Agriculture is the backbone of the economy, in addition to animal husbandry and tourism and 80% of the population in Spiti and Kinnaur are predominantly agro-pastoralists [36]. According to the 2011 census, the per capita annual income of the district is 2,289.57 USD (1,92,292 INR).

Ecologically, there is a robust community of small and large carnivores [41], including the vulnerable snow leopard (*Panthera uncia*). The two main prey species of the snow leopard – blue sheep *Pseudois nayaur* and ibex *Capra sibirica* – mostly occur in high altitude pastures and in relatively rugged terrain [4]. The livestock reared are sheep *Ovis aries*, goat *Capra hircus*, donkey *Equus asinus*, yak *Bos grunniens*, cattle *Bos indicus*, dzomo (a yak-cattle hybrid), and horses *Equus caballus*. The community has access to grazing pastures near the village over an area of approximately 70–100 *km*^2^ with traditional grazing and collection rights in the pastures, where cultivation is not permitted. Relatively smaller-bodied livestock such as goats, sheep, donkeys, cows and dzomo are taken grazing every day during the summer months while larger bodied livestock such as yak and horses are left to free-graze over most of the year except for a few weeks during extreme winter [38]. All livestock are stall fed during the winter months. Fodder is collected from the pastures and crop fields and stored as winter forage.

We used a multidisciplinary approach to study traditional production systems in the Indian-Trans Himalaya. We first used an ecological lens to perform field experiments comparing green pea, black pea, and barley on their growth performance. We then examined the genetic and nutritional diversity for black pea to assess how well these results align with farmers perceptions.

### 4.2 Field Experiment

The field study was conducted in March-September 2023 to evaluate the growth performance of three crop species: green peas, black peas, and barley under variable water treatments across altitudinal gradients. The experiment was carried at three sites characterized: low (4000m), mid (4200m), and high (4500m) elevations. The experiment was laid out in a randomized block design with a factorial arrangement, incorporating three factors: crop species, water treatments, and elevation levels. Each treatment combination was replicated three times, and thus a total of 81 plots were monitored. The seeds were sown uniformly across all plots and spacing was determined by interacting with farmers. The sowing density for green pea was lower than black pea seeds since black pea seeds are relatively smaller. The farmers also advised on sowing depth and bed preparation to ensure optimal growth conditions. The IRB eprotocol number is 71749.

We set up plots for green, black pea, and barley (1.5 by 1 meter) along a climate gradient in three villages (at elevations 4000m, 4200m, 4500m) with three water treatments. The water treatments represented varying levels of water availability: low, medium, and regular. The low water treatment simulated drought conditions, medium represented the less than standard recommended irrigation practice for the region, and regular simulated conditions of water used by farmers. The adjustment was made by altering the number of irrigation times rather than the quantity of water.

Throughout the cropping season, we monitored the plants for photosynthetic, non-reproductive and reproductive traits such as, germination rate, flowering time, height at maturity, leaf coloration (disease infestation), and flower/pod production. Data on growth parameters (plant height, leaf area, and biomass) were collected at regular intervals (every 7-10 days). Yield components were assessed at the time of harvest. Data were analyzed using ANOVA to determine the effects of the crop type, water treatment, elevation, and their interactions. Post hoc tests were performed to locate differences between treatment means at a significance level of p<0.05. All statistical analyses were carried out using R software.

### 4.3 Genetic diversity

For the genetic analysis, we first sampled fresh leaves and performed DNA extraction using DNeasy Plant Pro kit (Cat #69204). Illumina libraries were constructed using KAPA HyperPrep Kit and whole genome sequencing was performed at Medgenome labs (based out of Delhi, India) on an Illumina NovaSeq platform using a paired 150bp configuration. Illumina reads were mapped to the *P. sativum* reference genome (accession GCF_024323335.1) using BWA-MEM [30] and depth and breadth were calculated using SAMtools [31]. We want to note that the whole genome sequencing data generated for the black pea as part of this study has been deposited on NCBI - BioSample accession SAMN44380099.

We downloaded additional whole genome sequencing data from the recent paper [59] for P. sativum II, P. sativum III, P. sativum IV, and Pisum sativum var. arvense and also mapped these data to the reference genome using BWA-MEM. We list the whole genome sequencing data that was used for the PCA analyses (see Supplementary Information). The resultant alignment files (.bam files) were then run through ANSGD with the following flags “-uniqueOnly 1 -remove_bads 1 -only_proper_pairs 1 -trim 0 -C 50 -baq 1 -minMapQ 20 -minQ 20 -doCounts 1 -GL 1 -doMajorMinor 1 -doMaf 1 -skipTriallelic 1 -SNP_pval 1e-3 -doGeno 32 -doPost 1” to generate posterior probability of genotypes. We then used the ngsCover function from NGStools (Fumagalli et al. 2014) to generate a covariance matrix for each of the 7 chromosomes (2n=14) and visualised this data in a PCA using R.

### 4.4 Nutrition

The nutritional analysis for the black pea samples was done in collaboration with CSIR-Central Food Technological Research Institute (CFTRI) based out of Mysuru, India. The aim was to create a nutritional profile for black peas that includes the percentage per 100g of protein, carbohydrates, energy (kCal), and micronutrients such as zinc, magnesium, and iron.

## References

[1] Shahal Abbo, Simcha Lev-Yadun, and Avi Gopher. “Agricultural origins: centers and non-centers; a Near Eastern reappraisal”. Critical Reviews in Plant Science 29.5 (2010), pp. 317– 328.

[2] Eduardo Aguilera et al. “Agroecology for adaptation to climate change and resource depletion in the Mediterranean region. A review”. Agricultural Systems 181 (2020), p. 102809.

[3] Alexandre Antonelli. “Indigenous knowledge is key to sustainable food systems”. Nature 613.7943 (2023), pp. 239–242.

[4] Sumanta Bagchi and Charudutt Mishra. “Living with large carnivores: predation on livestock by the snow leopard (Uncia uncia)”. Journal of zoology 268.3 (2006), pp. 217–224.

[5] Martin Beniston. “Climatic change in mountain regions: a review of possible impacts”. Climatic change 59.1 (2003), pp. 5–31.

[6] Shriya Bhatt, Rashim Kumari, and Mahesh Gupta. “Development of soluble dietary fiber incorporated black pea protein slice: Physicochemical, textural, and rheological properties”. Measurement: Food 12 (2023), p. 100112.

[7] Teresa Borelli et al. “Local solutions for sustainable food systems: The contribution of orphan crops and wild edible species”. Agronomy 10.2 (2020), p. 231.

[8] Anwesha Borthakur and Pardeep Singh. “Indigenous knowledge systems in sustainable water conservation and management”. Water conservation and wastewater treatment in BRICS Nations. Elsevier, 2020, pp. 321–328.

[9] Eduardo S Brondízio et al. “Locally based, regionally manifested, and globally relevant: Indigenous and local knowledge, values, and practices for nature”. Annual Review of Environment and Resources 46.1 (2021), pp. 481–509.

[10] Pashupati Chaudhary and Kamaljit S Bawa. “Local perceptions of climate change validated by scientific evidence in the Himalayas”. Biology Letters 7.5 (2011), pp. 767–770.

[11] Aaron P Davis et al. “Lost and found: Coffea stenophylla and C. affinis, the forgotten coffee crop species of West Africa”. Frontiers in Plant Science 11 (2020), p. 521618.

[12] Fabrice AJ DeClerck et al. “Agricultural ecosystems and their services: the vanguard of sustainability?” Current opinion in environmental sustainability 23 (2016), pp. 92–99.

[13] Roy C Dudgeon and Fikret Berkes. “Local understandings of the land: Traditional ecological knowledge and indigenous knowledge”. Nature across cultures: Views of nature and the environment in non-Western cultures. Springer, 2003, pp. 75–96.

[14] FAO. “Indigenous Peoples’ food systems: Insights on sustainability and resilience from the front line of climate change”. Rome: Alliance of Biodiversity International, International Centre for Tropical Agriculture (CIAT) (2021).

[15] Joern Fischer, Tibor Hartel, and Tobias Kuemmerle. “Conservation policy in traditional farming landscapes”. Conservation letters 5.3 (2012), pp. 167–175.

[16] Michael-Shawn Fletcher et al. “Indigenous knowledge and the shackles of wilderness”. Proceedings of the National Academy of Sciences 118.40 (2021), e2022218118.

[17] Berhane Gebreslassie and Berhanu Abraha. “Distribution and productivity of Dekoko (Pisum sativum var. abyssinicum A. Braun) in Ethiopia”. Glob J Sci Front Res Biol Sci 16 (2016), pp. 45–57.

[18] Yirga Gufi et al. “Field pea diversity and its contribution to farmers’ livelihoods in northern Ethiopia”. Legume Science 4.4 (2022), e141.

[19] José Tomás Ibarra et al. “Mountain social-ecological resilience requires transdisciplinarity with Indigenous and local worldviews”. Trends in ecology & evolution (2023).

[20] Harman Jaggi, David Steinsaltz, and Shripad Tuljapurkar. “Temporal variability can promote migration between habitats”. Theoretical Population Biology 158 (2024), pp. 195– 205.

[21] Harman Jaggi et al. “Density dependence shapes life-history trade-offs in a food-limited population”. Ecology Letters 27.11 (2024), e14551.

[22] Devra I Jarvis et al. “An heuristic framework for identifying multiple ways of supporting the conservation and use of traditional crop varieties within the agricultural production system”. Critical Reviews in Plant Sciences 30.1-2 (2011), pp. 125–176.

[23] Runchun Jing et al. “The genetic diversity and evolution of field pea (Pisum) studied by high throughput retrotransposon based insertion polymorphism (RBIP) marker analysis”. BMC Evolutionary Biology 10 (2010), pp. 1–20.

[24] Santiago Kaulen-Luks et al. “Biocultural heritage construction and community-based tourism in an important indigenous agricultural heritage system of the southern Andes”. International Journal of Heritage Studies 28.10 (2022), pp. 1075–1090.

[25] Gemechu Keneni et al. “Genetic diversity for attributes of biological nitrogen fixation in Abyssinian field pea (Pisum sativum var. Abyssinicum) germplasm accessions”. Ethiopian Journal of Applied Science and Technology 4.2 (2013), pp. 1–20.

[26] Colin K Khoury et al. “Crop genetic erosion: understanding and responding to loss of crop diversity”. New Phytologist 233.1 (2022), pp. 84–118.

[27] Mayank Kohli et al. “Grazing and climate change have site-dependent interactive effects on vegetation in Asian montane rangelands”. Journal of Applied Ecology 58.3 (2021), pp. 539–549.

[28] Parviz Koohafkan and Miguel A Altieri. Globally important agricultural heritage systems: a legacy for the future. Food and Agriculture Organization of the United Nations Rome, 2011.

[29] OE Kosterin, VS Bogdanova, and AV Mglinets. “Wild pea (Pisum sativum L. subsp. elatius (Bieb.) Aschers. et Graebn. sl) at the periphery of its range: Zagros Mountains”. Vavilov Journal of Genetics and Breeding 24.1 (2020), p. 60.

[30] H Li. “Aligning sequence reads, clone sequences and assembly contigs with BWA-MEM”. arXiv preprint 1303.3997 (2013).

[31] Heng Li et al. “The sequence alignment/map format and SAMtools”. bioinformatics 25.16 (2009), pp. 2078–2079.

[32] Josh M Maurer et al. “Acceleration of ice loss across the Himalayas over the past 40 years”. Science advances 5.6 (2019), eaav7266.

[33] Charudutt Mishra, Herbert HT Prins, and Sipke E Van Wieren. “Diversity, risk mediation, and change in a Trans-Himalayan agropastoral system”. Human Ecology 31.4 (2003), pp. 595–609.

[34] Patricia Kefilwe Mogomotsi et al. “An analysis of communities’ attitudes toward wildlife and implications for wildlife sustainability”. Tropical Conservation Science 13 (2020), p. 1940082920915603.

[35] Sumit Mukherjee. “Changing Economy and Culture of Food in Spiti”. Food and Power: Expressions of Food-Politics in South Asia (2020), p. 1.

[36] Ranjini Murali, Stephen Redpath, and Charudutt Mishra. “The value of ecosystem services in the high altitude Spiti Valley, Indian Trans-Himalaya”. Ecosystem Services 28 (2017), pp. 115–123.

[37] Ranjini Murali et al. “Ecosystem service dependence in livestock and crop-based production systems in Asia’s high mountains”. Journal of Arid Environments 180 (2020), p. 104204.

[38] Ranjini Murali et al. “Indigenous governance structures for maintaining an ecosystem service in an agro-pastoral community in the Indian Trans Himalaya”. Ecosystems and People 18.1 (2022), pp. 303–314.

[39] Erez Naim-Feil et al. “Drought response and genetic diversity in Pisum fulvum, a wild relative of domesticated pea”. Crop Science 57.3 (2017), pp. 1145–1159.

[40] Rae Page et al. “Genome-wide association mapping of rust resistance in Aegilops longissima”. Frontiers in Plant Science 14 (2023), p. 1196486.

[41] J Patel et al. “Influence of predator suppression and prey availability on carnivore occurrence in western Himalaya”. Journal of Zoology 322.1 (2024), pp. 3–11.

[42] Ivette Perfecto and John Vandermeer. “Reflections on research agendas in agroecology: In search of a practical guide”. Journal of Agriculture, Food Systems, and Community Development 13.3 (2024), pp. 1–7.

[43] Katarina Perić et al. “The Role of Crop Wild Relatives and Landraces of Forage Legumes in Pre-Breeding as a Response to Climate Change”. Agronomy 14.7 (2024), p. 1385.

[44] Aghaghia Rahimzadeh. “Socio-economic and environmental implications of the decline of chilgoza pine nuts of Kinnaur, Western Himalaya”. Conservation and Society 18.4 (2020), pp. 315–326.

[45] Victoria Reyes-García et al. “Biocultural vulnerability exposes threats of culturally important species”. Proceedings of the National Academy of Sciences 120.2 (2023), e2217303120.

[46] Nicolas Rispail et al. “Genetic diversity and population structure of a wide Pisum spp. core collection”. International Journal of Molecular Sciences 24.3 (2023), p. 2470.

[47] M Santalla, Juana Marina Amurrio, and AM De Ron. “Food and feed potential breeding value of green, dry and vegetable pea germplasm”. Canadian Journal of plant science 81.4 (2001), pp. 601–610.

[48] Antonio Santoro et al. “A review of the role of forests and agroforestry systems in the FAO Globally Important Agricultural Heritage Systems (GIAHS) programme”. Forests 11.8 (2020), p. 860.

[49] Fausto O Sarmiento et al. “Applied montology using critical biogeography in the Andes”. Mountains: Physical, Human-Environmental, and Sociocultural Dynamics. Routledge, 2018, pp. 179–191.

[50] Astha Sharma and Mahesh Gupta. “Characterization of physicochemical, functional and antioxidant properties of western Himalayan black pea”. Journal of Agriculture and Food Research 13 (2023), p. 100607.

[51] RP Singh. “Integration and commercialization of local varieties under sub-optimal environments for food security, promoting sustainable agriculture and agro-biodiversity conservation”. MOJ Eco Environ Sci 3.2 (2018), pp. 65–67.

[52] Petr Smýkal et al. “Legume crops phylogeny and genetic diversity for science and breeding”. Critical Reviews in Plant Sciences 34.1-3 (2015), pp. 43–104.

[53] Pradeep Srivastava et al. “Late Pleistocene-Holocene morphosedimentary architecture, Spiti River, arid higher Himalaya”. International Journal of Earth Sciences 102 (2013), pp. 1967–1984.

[54] Kulbhushansingh Suryawanshi et al. “Estimating snow leopard and prey populations at large spatial scales”. Ecological Solutions and Evidence 2.4 (2021), e12115.

[55] Berhanu Abraha Tsegay and Berhane G Gebreegziabher. “Effects of terrains’ soil and altitude on performance of Abyssinian pea (Pisum sativum var. abyssinicum A. Braun) landraces of Ethiopia”. Biodiversitas Journal of Biological Diversity 20.12 (2019).

[56] Shripad Tuljapurkar et al. “From disturbances to nonlinear fitness and back”. bioRxiv (2023), pp. 2023–10.

[57] Rahul Verma. “The revival of ancient grains: Millets”. International Journal of Political Science and Governance (2023).

[58] Alexander Wezel et al. “Agroecological principles and elements and their implications for transitioning to sustainable food systems. A review”. Agronomy for Sustainable Development 40 (2020), pp. 1–13.

[59] Tao Yang et al. “Improved pea reference genome and pan-genome highlight genomic features and evolutionary characteristics”. Nature genetics 54.10 (2022), pp. 1553–1563.

